# Evidence-driven biases in alternative splicing inferred from NCBI Eukaryotic Genome Annotation Pipeline metadata

**DOI:** 10.1101/2025.05.27.656353

**Authors:** Rebeca de la Fuente, Wladimiro Díaz-Villanueva, Vicente Arnau, Andrés Moya

## Abstract

The NCBI Eukaryotic Genome Annotation Pipeline (EGAP) predicts coding sequences by integrating transcriptomic and proteomic data with computational approaches, providing structural information that can be used to infer alternative transcript isoforms. However, accurate estimation of alternative splicing events depends on high-quality genome annotations, particularly in genome-wide analyses. The extent to which annotation pipelines influence these inferred splicing patterns has remained largely unexplored. In this study, we quantify potential biases associated with the EGAP annotation pipeline, and find that specific annotation features strongly influence these estimates, particularly the percentage of coding sequences that are supported by experimental evidence. Further, we implemented a polynomial regression model to normalize splicing levels, generating an adjusted metric that minimizes evidence-driven biases. This framework may serve as a basis for future investigations into splicing complexity and comparative genomics.

## Introduction

Alternative splicing is a fundamental mechanism that expands transcriptomic and proteomic diversity in eukaryotes by generating multiple transcript isoforms from a single gene through differential exon usage [Nilsen and Graveley, 2010, Barbosa-Morais et al., 2012, Mattick, 2009, Grosso et al., 2008]. It plays a central role in regulating critical biological processes, including immune system function, neuronal development, and cellular homeostasis [Shalek et al., 2013, Weyn-Vanhentenryck et al., 2018, Tapial et al., 2017]. By expanding the functional repertoire of the genome, alternative splicing allows for the fine-tuned regulation of complex processes like immune system signaling [Lynch, 2004, Martinez and Lynch, 2012]. In the nervous system, alternative splicing has been shown to be essential for neuronal differentiation, synapse formation, and plasticity, providing molecular diversity required for the intricate architecture and functionality of the brain [Li et al., 2007, Raj and Blencowe, 2015]. Moreover, dysregulation of splicing is increasingly recognized as a major contributor to a wide range of diseases, including cancer, autoimmune disorders, and neurodegenerative conditions [Scotti and Swanson, 2016, Wang and Cooper, 2007]. Furthermore, it is involved in differentiation processes across multiple levels of biological organization, including tissue differentiation, the emergence of distinct morphs, and speciation events [Lyko et al., 2011, Harr and Turner, 2010, Steward et al., 2022, Grantham and Brisson, 2018]. Yet, our understanding of how alternative splicing contributes to diversity across the eukaryotic tree of life is still limited. To date, most research has been conducted in a narrow set of well-studied model organisms, such as *Homo sapiens, Drosophila melanogaster*, and *Arabidopsis thaliana* [Pan et al., 2008, Wang et al., 2008, Graveley et al., 2011, Marquez et al., 2012]. Barbosa-Morais et al. [2012] and Tilgner et al. [2015] demonstrated that, although the human genome reference improves annotation accuracy, it may fail to capture the full spectrum of splicing diversity in non-model organisms. They found that within just six million years, the splicing profiles of physiologically equivalent organs have evolved to be more closely associated with the identity of a species than with the organ type. Furthermore, most studies predominantly focused on specific genes, tissues, or developmental stages, often under tightly controlled laboratory conditions. As a result, the current understanding of its regulation and function is largely context-specific, and its generalizability across taxa remains limited [Steijger et al., 2013, Merkin et al., 2012, Tress et al., 2017]. As a consequence of differences in scientific goals and resource availability, different studies employ distinct sequencing technologies (e.g., short-read vs. long-read RNA-Seq), library preparation protocols, and computational pipelines for transcript assembly, isoform quantification, and genome annotation [Steijger et al., 2013, Tardaguila et al., 2018]. In particular, the use of specific annotation frameworks has led to uneven transcript coverage and varying sensitivity for detecting low-abundance or tissue-specific isoforms [Tress et al., 2017, Zhang et al., 2020]. This methodological heterogeneity introduces systematic biases in cross-study analyses, ultimately limiting our ability to generalize splicing patterns at the species level and assess their evolutionary significance.

In response to these limitations, the scientific community has made increasing efforts to create unified annotation resources and cross-species frameworks. Initiatives such as GENCODE [Frankish et al., 2018], Ensembl [Howe et al., 2020], and RefSeq [O’Leary et al., 2015] have progressively integrated manual curation and multiple sources of experimental evidence to enhance transcriptome completeness and reduce annotation inconsistencies. Also, the adoption of shared gene models across platforms and the inclusion of long-read sequencing data have improved isoform resolution and supported the detection of tissue-specific transcripts even in less-studied organisms [Martin and Wang, 2011, Liu et al., 2023]. The Eukaryotic Genome Annotation Pipeline (EGAP) integrates evidence from RNA-Seq, expressed sequence tags (ESTs), and protein alignments, alongside computational predictions generated by tools such as Gnomon [Thibaud-Nissen et al., 2013]. Gnomon is a gene prediction program developed by the National Center for Biotechnology Information (NCBI) that integrates experimental evidence with computational ab initio methods. It employs advanced algorithms, including Hidden Markov Models, to predict coding regions. This dual approach ensures accurate predictions even in cases where experimental evidence is limited or unavailable. A key feature of Gnomon is its ability to handle complex genomic data through its Chainer algorithm, which assembles overlapping alignments into coherent transcripts. It also accounts for alternative splicing, producing multiple transcript variants for a single gene when supported by the evidence. Furthermore, functional information is enriched through tools like InterProScan, which assigns Gene Ontology (GO) terms and other functional annotations to protein-coding genes [Blum et al., 2024]. The output of Gnomon includes predicted transcripts and proteins, which are assigned to unique RefSeq accessions (e.g., XM, XR, XP prefixes) to distinguish them from known sequences. The result is a comprehensive and non-redundant set of genomic features, extensively annotated with structural and functional information, and enriched with cross-references to various genomic and functional annotation resources. Despite these advancements, a critical gap remains in our understanding of how annotation pipelines systematically influence transcriptome-based measurements, such as estimates of alternative splicing levels.

Alternative splicing levels can be inferred from the structure and redundancy of annotated coding sequences (CDSs), which can be computed from the annotation files provided by the NCBI EGAP. This approach relies on identifying the number of distinct coding sequences mapped to a given genomic locus, which can be interpreted as a proxy for transcript isoform diversity. By aggregating this information at the genome level, it is possible to derive quantitative splicing metrics such as the *Alternative Splicing Ratio* (ASR), a metric that was previously proposed in a comparative study involving 1,494 species spanning the entire tree of life [de la Fuente et al., 2025]. Given the wide use of EGAP for annotating RefSeq genomes, it is essential to identify which metadata variables and methodological components introduce the greatest biases in inferred splicing complexity. Quantifying these sources of bias is a prerequisite for building correction models that allow for meaningful comparisons across taxa. In addition to the annotation files, each genome processed through EGAP is accompanied by a detailed annotation report that provides extensive metadata on the source and type of evidence used for each predicted feature. These reports include variables such as the number of CDSs supported by transcript or protein alignments, the fraction of CDSs derived from ab initio prediction, and other quality indicators relevant to transcript annotation. This metadata represents a valuable but underexplored resource for quantifying annotation-derived biases and evaluating their impact on downstream transcriptomic analyses.

The pipeline prioritizes curated RefSeq transcripts and genomic sequences when available, using them to directly annotate CDSs or to guide predictions. However, most annotations are based on short-read RNA-Seq data, which cannot provide information about the exact transcript being expressed in a sample. Addressing this requires long-read sequencing, which is used in the EGAP but not extensively. Additionally, mappings from long reads are not reported in the annotation files, and while individual features include counts of supporting samples, the reports do not provide a detailed list of the samples associated with each feature. For regions lacking direct evidence, predictions are supplemented by ab initio methods that leverage genomic signals like start codons, stop codons, and splice sites. For example, for alignments with partial cDNAs or RNA-Seq data without complete coverage, coding regions are predicted using the Gnomon model. All cDNAs with coding sequence scores above a specific threshold are classified as coding, ensuring that only sequences with strong evidence are included as CDSs in the annotation. This threshold helps filter out non-coding regions and low-confidence predictions, improving the overall accuracy of annotations. The integration of experimental data and computational predictions ensures that the annotated CDSs are as accurate and comprehensive as possible, but the quality and depth of the input data, such as RNA-Seq, can significantly influence the final results. High read depth enhances the ability to identify low-abundance isoforms and validate splicing events across complex loci, as noted by Yang et al. [2024]. In particular, differences in sequencing depth across genome annotations may overrepresent highly expressed isoforms while masking rare or condition-specific splicing events. Tissue diversity in RNA-Seq datasets is another critical factor, as it contributes to transcriptomic complexity and functional diversity. Wang et al. [2008] emphasized that incorporating data from diverse tissues is crucial for revealing splicing patterns specific to certain biological contexts. The number of experimental runs further complements splicing analyses by introducing biological and technical replicates, which increase statistical power and the robustness of results. Steijger et al. [2013] highlighted that balanced experimental designs with multiple runs per condition are essential for reliable detection of splicing events across conditions and replicates. In addition to biases related to CDS annotation depth, alignment strategies rely heavily on the quality of the reference genome and the capacity of aligners to handle intronic regions. In turn, the quality of genome assembly can influence the detection of splice variants [Liu et al., 2023]. Furthermore, sequencing errors can introduce false splice sites, leading to incorrect identification of splicing events [Sturgill et al., 2013, Kurmangaliyev and Gelfand, 2008]. Recognizing these challenges, some studies have developed de novo approaches to improve splicing analysis without relying on a reference genome. Relevant to this field, we find the tool KisSplice, which identifies splicing events directly from RNA-seq data [Sacomoto et al., 2012], and SOAPdenovo-Trans, which focuses on reconstructing full-length transcripts to enhance isoform detection [Xie et al., 2014]. These methods mitigate the limitations of incomplete or low-quality assemblies. However, while de novo assembly can identify novel isoforms, it is computationally intensive and prone to errors in regions with low coverage [Martin and Wang, 2011]. These challenges can be mitigated by using high-quality assemblies with long contigs or scaffolds. Moreover, high-quality assemblies significantly minimize sequencing errors and facilitate the effective use of strand-specific RNA-Seq protocols, which are essential for resolving overlapping transcripts, particularly in highly spliced genomes.

In this study, we estimate alternative splicing levels from the genome annotation files provided by the NCBI annotation files, and extract detailed features from the corresponding annotation reports to evaluate evidence-driven biases that may affect these estimates. In addition to biases related to the depth and quality of CDS annotations, our analysis incorporates genome assembly statistics, tissue diversity, and other relevant features to detect potential distortions in transcript-level metrics. By combining this information with genome-wide annotations across hundreds of eukaryotic species, we identify the most influential sources of systematic error. Furthermore, we introduce a normalization procedure designed to correct annotation-driven deviations in splicing estimates, enabling more reliable cross-species comparisons and supporting downstream analyses of transcriptomic complexity.

## Materials and Methods

### Data Collection and Processing

We collected a dataset of annotated eukaryotic genomes through the NCBI, focusing specifically on assemblies curated by the RefSeq project [Sayers et al., 2023]. Genome and annotation metadata were retrieved from the NCBI FTP service, which serves as the official distribution platform for high-quality genomic resources [Site, 2024]. These data are organized in a taxonomically structured directory and include both assembly-level statistics and detailed annotation reports. While assembly reports are provided as text files, annotation reports are available as XML files on the genomes FTP site. For detailed information, see Data Availability section.

To ensure data reliability, we applied three filtering criteria. First, we restricted our dataset to assemblies annotated at the chromosome or complete genome level, which correspond to the highest-confidence genome builds. Second, we restricted the dataset to assemblies from the RefSeq database that have been processed through the EGAP annotation pipeline, thereby ensuring consistency in annotation methodology across species. Third, we focused exclusively on multicellular eukaryotes and grouped them into five major taxonomic categories: mammals (133), birds (77), fish (169), arthropods (187), and plants (128). The resulting dataset included 694 species classified within distinct multicellular eukaryotic clades. Although the alternative splicing ratio was computed for all 694 species using their corresponding genome annotation files, annotation reports were only available for a subset of 670 species. As a consequence, the analysis of variables potentially influencing alternative splicing estimates was restricted to those species for which complete annotation metadata was successfully retrieved. Taxonomic assignments were established based on a phylogenetic representation generated with the Interactive Tree of Life (iTOL) tool [Kumar et al., 2022], and subsequently validated using the NCBI Taxonomy database [Federhen, 2011, Sayers et al., 2023]. Based on the annotation-derived metadata and assembly reports, we extracted a total of 23 relevant variables that served as the foundation for evaluating annotation quality and identifying systematic biases in alternative splicing estimates.

### Alternative Splicing Ratio

A novel genome-scale metric of alternative splicing was proposed in de la Fuente et al. [2025] for the comparative study of splicing patterns across the tree of life. This metric allowed large-scale evolutionary analyses of transcript diversity based on gene annotation data. We refer to this metric as the *Alternative Splicing Ratio* (ASR), which measures the degree of transcriptomic diversity by dividing the total number of annotated coding transcripts by the number of protein-coding sequences in a given genome. While the metric can be calculated at the level of individual genes, it is generalized to the whole genome by projecting all annotated coding sequences from transcripts onto their corresponding genomic positions. Condensing this information into a single numerical index, it serves as a standardized metric to compare alternative splicing levels across species.

The alternative splicing ratio is computed from the RefSeq GFF annotation files. For each species, we parsed the corresponding annotation file and extracted information about all protein-coding genes and their transcript isoforms. The files are organized using a parent-child structure, where genes act as parent elements to their corresponding transcripts (mRNAs), and each transcript is further linked to its CDSs. This hierarchical format reflects the biological relationship between a gene and its alternative splicing isoforms, with each mRNA representing a distinct transcript variant. The CDSs associated with each mRNA define the protein-coding regions retained after splicing, ultimately specifying a unique protein isoform. In our analysis, we focused exclusively on CDSs with clearly defined hierarchical links, excluding pseudogenes and duplicated genes. The data is structured as tab-separated tables, where each row corresponds to a genomic feature—such as a gene, mRNA, or CDS—and each column contains specific attributes, including unique identifiers, genomic positions, and functional annotations. For each genomic feature, we extracted its start and end coordinates, which allowed us to calculate the length of individual regions, such as genes, transcripts, and coding sequences. Thus, the total gene content was quantified as the cumulative number of base pairs that fall within annotated gene intervals. Similarly, coding size was computed by summing the nucleotides in the genome annotated as CDS. In contrast, the transcript domain is defined as the total number of nucleotides annotated as CDS, but in this case, each transcript was considered independently—that is, the CDSs of all isoforms were summed without collapsing overlapping regions. The transcript domain reflects the total coding potential at the transcript level and accounts for redundancy across isoforms. Thus, the ASR was computed as the ratio between the total size of the transcript domain—defined as the cumulative length of all CDSs across all annotated transcripts—and the coding size, which corresponds to the projection of these CDSs onto the genome, counting each nucleotide only once regardless of how many isoforms share it. This comparison captures the extent to which coding regions are reused across different transcripts, providing a genome-wide estimate of alternative splicing activity.

### Statistical Analyses

We performed Spearman correlation analyses using R, aiming to quantify the pairwise associations between the alternative splicing ratio and annotation-related variables. Spearman’s method was chosen because it captures monotonic relationships, allowing us to detect associations that are not strictly linear. The resulting correlation matrix between variables provided a basis for subsequent multivariate modeling and variable selection.

We also performed a multivariate analysis to explore the joint contribution of annotation variables to alternative splicing ratio. To reduce multicollinearity and avoid redundancy among highly correlated predictors, we applied a Least Absolute Shrinkage and Selection Operator (LASSO) regression, a regularized linear modeling approach particularly effective for high-dimensional data and variable selection [Tibshirani, 2018]. LASSO introduces a penalty term controlled by the regularization parameter *λ*, which shrinks the absolute values of regression coefficients, setting many to exactly zero. This allows the model to retain only the most informative predictors while preventing overfitting. Importantly, LASSO evaluates all variables simultaneously, and its output is not influenced by the order in which predictors are entered into the model. To implement this analysis, we first split the dataset into a training set (80%) and a test set (20%). LASSO regression was then fitted to the training data using the *glmnet* package in R [Friedman et al., 2010]. To identify the optimal level of penalization, we performed k-fold cross-validation, computing the mean squared error across a range of *λ* values. Finally, the performance of the trained model was evaluated on the test set.

## Results

### Annotation-Related Metrics

Tables 1 to 3 provide a conceptual description of the 23 metrics extracted from the NCBI annotation reports and considered in our evaluation of genome annotation quality. In Table 1, we summarize the set of metrics used to evaluate genome assembly quality. Scaffold and contig counts, together with their corresponding N50 values, reflect the degree of assembly fragmentation. Higher N50 values and lower scaffold or contig counts indicate a more contiguous assembly, which is critical for accurate genome annotation. Gap length is also considered, as it serves as an indicator of assembly continuity, and has been previously recognized as a robust measure of assembly quality. Table 2 summarizes the set of metrics corresponding to the molecular datasets that support the annotation process. This includes the number of protein and transcript sequences retrieved from the Entrez database, which are mapped to the genome assembly being annotated and used by Gnomon to gene prediction. These sequences include mRNAs, ESTs, and curated protein records from RefSeq and other sources. Transcript sequences are aligned using Splign, while protein sequences are aligned using ProSplign, both tools developed by the NCBI. The resulting alignments serve as evidence to support gene prediction. Gnomon, the ab initio gene prediction algorithm employed by EGAP, combines this external evidence with computational modeling to generate gene structures, including exon-intron boundaries and CDS annotations. When high-confidence alignments occur, Gnomon uses them for predictions, prioritizing models that match the experimental data. In regions lacking strong evidence, Gnomon relies more heavily on intrinsic sequence signals to infer gene structure. Table 2 also includes metrics related to RNA-Seq data attributes, such as the total number of reads, tissue diversity, and the number of experimental runs. Collectively, these variables serve as quantitative indicators of the extent of experimental support and annotation depth, allowing us to assess their influence on the genome annotation process. Finally, Table 3 summarizes the metrics related to the annotated CDSs, including those supported by experimental evidence and those inferred through computational prediction. Analyzing the impact of these metrics on alternative splicing levels will allow us to better understand the degree of reliance on predictive modeling and the balance between evidence-based and computational approaches.

**Table 1.** Summary of annotation metadata metrics derived from the NCBI annotation reports. These metrics describe assembly-associated metadata, including scaffold and contig statistics, N50 values, and gap-related information. The metric for both Scaffold N50 and Contig N50, as well as the gap length, is measured in base pairs (bp).

| Metric | Description |
| --- | --- |
| Number of Gaps | Total number of gaps present in the genome assembly |
| Scaffold Count | Total number of scaffolds in the assembly |
| Scaffold N50 | The length of the smallest scaffold such that 50% of the total genome length is covered by scaffolds of this length or longer |
| Contig Count | Total number of contigs in the assembly |
| Contig N50 | The length of the smallest contig such that 50% of the total genome length is covered by contigs of this length or longer |
| Gap Length | Total length of gaps in the genome assembly |
| Gap Length (%) | The percentage of the total genome length that consists of gaps |

**Table 2.** Summary of annotation metadata metrics associated with the molecular data supporting the annotation process. The table includes the number of proteins and transcripts retrieved from the Entrez database, and RNA-Seq data characteristics, including read counts, tissue diversity, and experimental runs.

| Metric | Description |
| --- | --- |
| Number of proteins | Total number of protein sequences retrieved from the Entrez database for the annotation process |
| Number of transcripts | Total number of transcript sequences retrieved from the Entrez database for the annotation process, including mRNA, ESTs, and other RNA sequences |
| Number of proteins from *H. Sapiens* | Total number of protein sequences originating from *Homo Sapiens* and retrieved from the Entrez database |
| Number of transcripts from *H. Sapiens* | Total number of transcript sequences originating from *Homo Sapiens* and retrieved from the Entrez database |
| Number of RNA-Seq reads | Total count of RNA-Seq reads retrieved from the Sequence Read Archive that were utilized during the annotation process to support gene prediction |
| Number of tissues | The number of distinct tissue types represented in the RNA-Seq data |
| Number of runs | Total count of individual RNA-Seq experimental datasets (runs), each corresponding to a specific biological sample, condition, or replicate |

**Table 3.** Summary of annotation metadata metrics describing the number of CDSs generated during the annotation process, including total counts, those derived from computational models, and those supported by experimental evidence.

| Metric | Description |
| --- | --- |
| **CDSs** | Total number of CDSs annotated in the genome |
| **CDSs from model** | Total number of CDSs predicted by computational models, such as ab initio methods or guided by evidence but not experimentally validated |
| **Fully supported CDSs** | Total number of CDSs supported by direct experimental evidence, such as RNA-Seq or protein alignments |
| **Known CDSs** | Number of CDSs that correspond to sequences already present in curated databases like RefSeq. These are derived from previously validated annotations |
| **CDSs with > 5% ab initio** | Total number of CDSs where more than 5% of their bases are annotated using ab initio methods. This indicates regions where computational predictions were necessary due to insufficient experimental evidence |
| **CDSs from model (%)** | The percentage of total CDSs that are derived from computational models |
| **Known CDS (%)** | The percentage of total CDSs that are known from curated databases |
| **Fully supported CDSs (%)** | The percentage of total CDSs supported by experimental evidence |
| **CDSs with > 5% ab initio (%)** | The percentage of total CDSs where more than 5% of their bases rely on ab initio predictions |

### Associations Between Annotation Metadata and Alternative Splicing

In this section, we explore how the previously described set of 23 annotation-related quality variables may influence the quantification of alternative splicing levels through correlation analyses. In the first block of Table 4 and Fig. 1 of the Supplementary Material, we observe weak correlations between the alternative splicing ratio and assembly-related metrics, with the strongest correlation found with Contig N50. These results suggest that more fragmented assemblies may slightly reduce the number of alternative isoforms detected, but the weakness of the correlations indicates that it may not play a major role in shaping alternative splicing estimates. Although a relationship may be masked by noise or the influence of additional variables, the overall lack of strong correlations suggests that assembly quality is not a major determinant of the predicted levels of alternative splicing. We also evaluate the interdependencies among assembly metrics. The number of gaps shows a strong positive correlation with gap length and a strong negative correlation with contig N50. In turn, total gap length shows a strong association with its proportion relative to genome size. Finally, a positive correlation is observed between scaffold and contig counts, which suggests that assemblies with many scaffolds are also composed of numerous small contigs, a hallmark of low contiguity. Altogether, assembly quality can be evaluated using indicators such as gap count and scaffold continuity metrics like Scaffold N50.

**Table 4.**
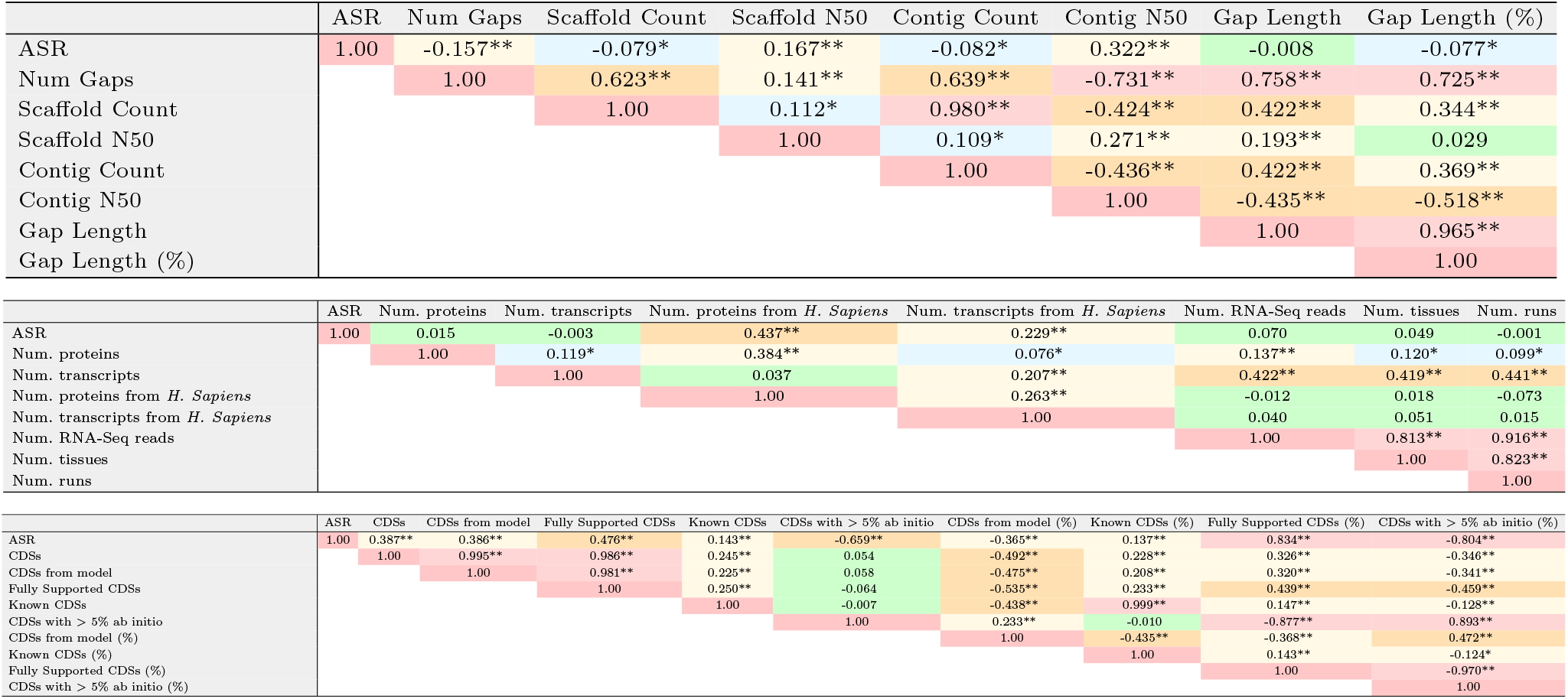
Pairwise Spearman correlation matrices between the alternative splicing ratio and the annotation-related metrics described in Tables 1–3. The first subtable displays correlations between ASR and assembly quality metrics (Table 1); the second includes correlations with experimental support metrics (Table 2); and the third corresponds to CDS-related annotation output metrics (Table 3). Asterisks indicate significance levels: *p <* 0.05 (*), *p <* 0.001 (**). Cell colors reflect the strength of the correlation coefficients.

**Fig. 1.**
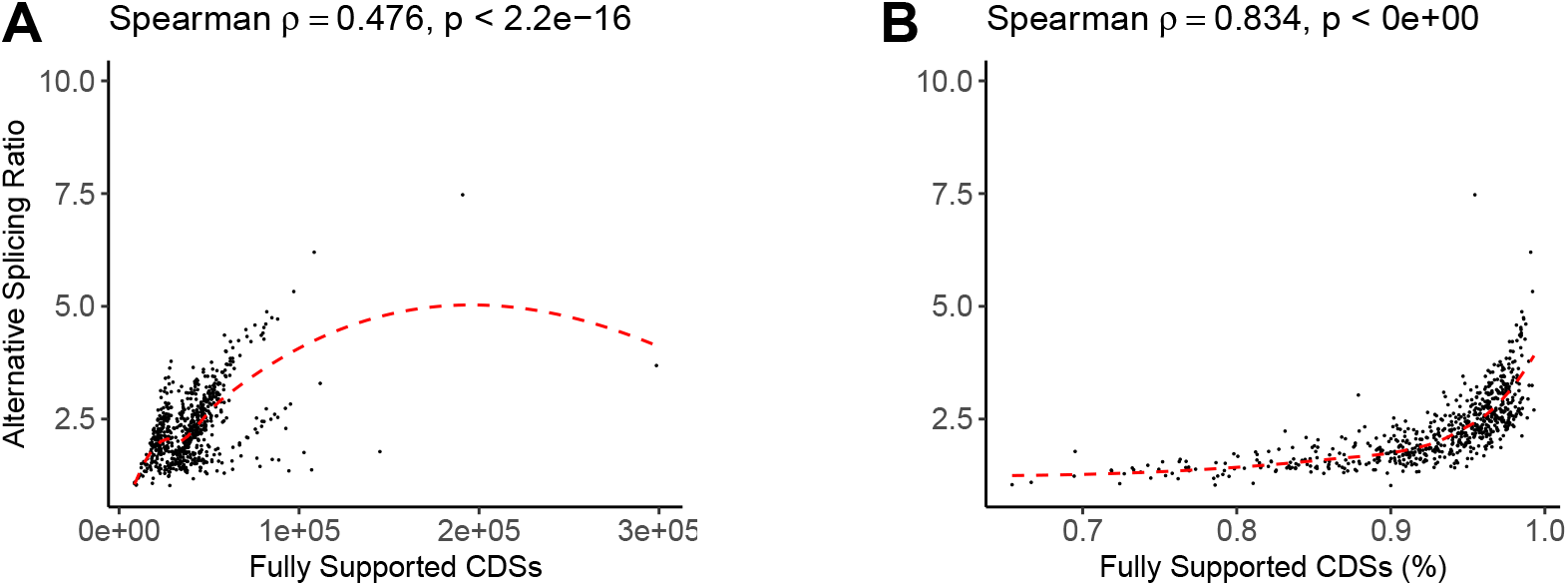
Spearman correlation between ASR and annotation metrics related to CDSs. (A) Correlation between ASR and the absolute number of fully supported CDSs. (B) Correlation between ASR and the percentage of fully supported CDSs. These variables, which are among the most relevant predictors of splicing estimates, are described in detail in Table 3.

Next, the intermediate block in Table 4 shows pairwise associations involving the experimental evidence variables listed in Table 2 (see also Fig. 2 of Supplementary Material). Alternative splicing levels do not correlate with the total number of proteins and transcripts retrieved from the Entrez database. However, they do show weak but significant correlations with the subset of evidence derived from *Homo sapiens*. Specifically, both protein and transcript datasets from this species exhibit positive correlations with the alternative splicing ratio, suggesting that the abundance of human-derived data may slightly inflate the observed splicing levels. However, it is important to note that these correlations are very weak, and no significant associations were found with the remaining variables, namely the number of RNA-Seq reads, the diversity of tissues, or the number of runs. These results suggest that, while the overall amount of experimental evidence does not appear to influence alternative splicing estimates, a slight inflation may occur in annotations with a disproportionate representation of data derived from *Homo sapiens*. We now examine the relationships among the evidence-derived variables. We observe strong correlations among the number of RNA-Seq reads, tissue diversity, and the number of runs. This reflects the structure of typical RNA-Seq-based datasets and the inherent interdependence among experimental design variables, where increased tissue diversity is accompanied by a higher number of sequencing runs and greater sequencing depth. Beyond these correlations, the number of transcripts is the only variable showing any additional association with these factors; however, the correlation is very weak, and none of the other variables showed relevant correlations.

**Fig. 2.**
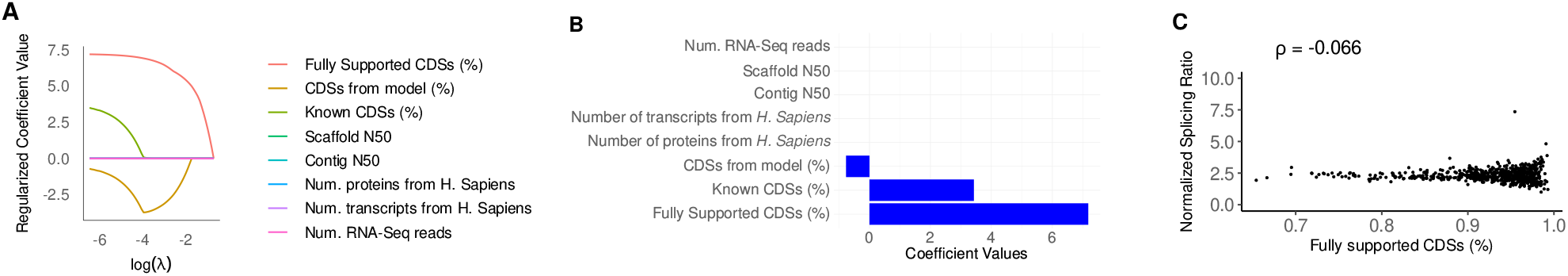
LASSO regression results for predicting alternative splicing ratio (ASR). (A) Evolution of model coefficients as the regularization parameter (*λ*) increases, illustrating how less informative variables are progressively excluded from the model. Only three predictors remain with non-zero coefficients at high penalization levels: Fully Supported CDSs (%), Known CDSs (%), and CDSs from Model (%). (B) Final coefficient values using the optimal penalty of *λ*, which achieves a balance between model simplicity and predictive power. (C) Spearman correlation between the normalized splicing ratio (ASR^*∗*^) and the percentage of fully supported CDSs, which shows a non-significant association after normalization.

Whereas variables describing the amount of experimental evidence show an overall lack of correlation with the alternative splicing ratio, we observe consistent correlations with all variables related to annotated CDSs, highlighting the potential influence of the annotation process in shaping observed splicing complexity. In the final section of Table 4 and Fig. 3 of Supplementary Material, we observe a variety of correlations between the alternative splicing ratio and the CDS-related annotation variables, spanning from very weak to strong. First, very weak correlations are found with both the number of known CDSs and the proportion of known CDSs. Weak correlations are observed with the total number of CDSs, the number of CDSs derived from computational models, and the proportion of CDSs annotated from models. Next, moderate correlations are detected for the number of fully supported CDSs and for the number of CDSs with more than 5% of their bases annotated via ab initio methods. Finally, strong correlations are observed with the percentage of fully supported CDSs and with the proportion of CDSs containing more than 5% ab initio content. These findings indicate that it is not merely the volume of data that shapes splicing estimates, but how effectively that evidence is integrated into the annotation process. In this context, Fully Supported CDS (%) emerges as a proxy for the degree to which empirical information is translated into annotated isoform diversity. Thus, the degree of evidence incorporated during transcript annotation, rather than the raw input data, appears to be the main factor shaping observed levels of alternative splicing.

We also observe very strong correlations among several of the CDS-related annotation variables, indicating that these metrics are closely interdependent. The number of known CDSs shows a very strong correlation with the proportion of known CDSs. Similarly, the total number of CDSs, the number of fully supported CDSs, and the number of CDSs predicted by computational models are all strongly correlated with one another. These strong correlations suggest that CDS-related variables do not represent independent annotation categories, but rather interrelated components of the overall annotation process. Genomes with more CDSs tend to exhibit higher counts of both model-predicted and fully supported CDSs, reflecting an underlying dependency driven by annotation depth and computational inference. In practice, the annotation pipeline—particularly Gnomon— integrates both evidence-supported and model-derived CDSs. A single CDS may be partially constructed based on ab initio predictions and partially informed by transcript or protein alignments. In this sense, model predictions are frequently guided by evidence, meaning that computationally derived annotations incorporate empirical support. As a result, fully supported CDSs and those obtained through modeling are not mutually exclusive, but instead reflect overlapping annotation strategies. This highlights the importance of interpreting these metrics as interdependent components of a unified annotation framework. While absolute counts of CDSs—whether model-derived or evidence-supported—reflect overall annotation depth, the relative proportions of these categories offer complementary information. In particular, we observe strong correlations among Fully Supported CDSs (%), CDSs with more than 5% ab initio, and their corresponding proportion, yet these do not correlate with the absolute number of fully supported CDSs. This suggests that the annotation strategy, in terms of reliance on evidence versus ab initio prediction, is better captured by relative metrics than by raw counts, and may play a more direct role in shaping transcript-level features such as splicing diversity.

As illustrated in Figure 1, the correlation between the alternative splicing ratio and the absolute number of fully supported CDSs is noisy and suggests the presence of multiple regimes across species. This dispersion likely reflects underlying heterogeneity in genome size and gene content, which may introduce variability in how annotation depth translates into splicing detection. In contrast, when alternative splicing values are plotted against the percentage of fully supported CDSs, the relationship becomes much more consistent and exhibits a smooth, non-linear trend with minimal scatter. This contrast implies that it is not the absolute amount of empirical evidence that best explains observed levels of alternative splicing, but rather its relative abundance within the total annotation set. Thus, the percentage of fully supported CDSs normalizes for other confounding factors and captures the degree to which empirical data shapes isoform diversity. These findings reinforce the view that annotation-based metrics like ASR are intrinsically shaped by the architecture of the annotation process, emphasizing the importance of normalization strategies to mitigate annotation-driven biases.

We next conducted a multivariate analysis to identify the most influential predictors of alternative splicing (see Methods). To this end, we applied a LASSO regression, a regularization method that selects a minimal subset of variables while penalizing model complexity. This approach enabled us to detect complex dependencies among variables that may not be evident when examined individually. By considering all variables simultaneously, the analysis revealed which factors most strongly contribute to variation in splicing levels. Based on the interdependencies identified in previous analyses, we selected a subset of eight representative variables for the multivariate analysis. This selection aimed to retain the relevant predictors while minimizing collinearity and redundancy. The variables included were: Contig N50, Scaffold N50, number of proteins from *Homo sapiens*, number of transcripts from *Homo sapiens*, number of RNA-Seq reads, Fully supported CDSs (%), model-derived CDSs (%), and known CDSs (%).

In Figure 2(A), we illustrate how the LASSO model evolves as the regularization parameter (*λ*) increases. At lower *λ* values, the model includes a greater number of predictors, some of which may contribute only marginally to the explanation of splicing variability. As *λ* increases, the model imposes stronger penalties on less informative variables, progressively shrinking their coefficients toward zero. This results in a more compact model that retains only the most relevant predictors. In our case, we found only three key predictors: fully Supported CDSs (%), Known CDSs (%), and CDSs derived from models (%). These variables remain relevant for predicting alternative splicing even under strong penalization, while the remaining predictors are excluded from the model altogether. As a consequence, although many variables are correlated with splicing levels, only a few provide independent, non-redundant information.

In the pairwise correlation analyses, the percentage of CDSs from the model and the percentage of known CDSs showed only weak correlations with ASR. Interestingly, despite their weak pairwise associations, these two variables emerged as among the most predictive in the multivariate model. This indicates that they may capture complementary aspects of the annotation process that become informative when considered in combination with other variables. In contrast, we found that the percentage of fully supported CDSs showed a very strong correlation with ASR values. Consistently, this variable also emerged as the most predictive in the multivariate model, indicating that the evidence support of annotated CDSs may systematically bias ASR estimates. Species with higher proportions of fully supported CDSs tend to show elevated levels of alternative splicing, potentially reflecting differences in annotation quality rather than true biological variation.

Using the optimal level of penalization determined during model fitting (*λ*), we estimated the contribution of each annotation feature to the prediction of ASR values (Figure 2B). The final model retained only the three annotation variables previously identified, all with non-zero coefficients. Among them, the percentage of fully supported CDSs had the highest coefficient, highlighting it as the most influential variable in the model. All other variables had coefficients near zero, indicating a negligible impact on the model. Finally, we evaluated how well the model could predict ASR values in a separate dataset that was not used during model training. The results showed a good level of predictive accuracy, with the model explaining approximately 60% of the variation in ASR values. This indicates that the selected annotation variables capture a substantial portion of the differences in alternative splicing observed across species.

Since the percentage of fully supported CDSs was identified as the main driver of annotation-related biases in ASR, we normalized ASR values based on its relationship with this variable. This approach allowed us to account for variation that may stem from differences in annotation quality. To capture the non-linear pattern of this association, which is illustrated in Figure 1B, we fitted a fourth-degree polynomial regression model. The model provided a good fit to the data, explaining approximately 60% of the variability in ASR values (multiple *R*^2^ = 0.605, and adjusted *R*^2^ = 0.602). The estimated coefficients for all polynomial terms (up to the fourth degree), along with their statistical significance, are presented in Table 5. Thus, we normalized ASR values according to the following formulation:

**Table 5.** Coefficients of the fourth-degree polynomial regression model used to describe the relationship between ASR and the percentage of fully supported CDSs. The table includes the estimated value, standard error, t-value, and p-value for each polynomial term. All terms are statistically significant, indicating that higher-order components contribute meaningfully to capturing the non-linear association.

| Term | Estimate | Std. Error | t-value | p-value |
| --- | --- | --- | --- | --- |
| Intercept | 2.3180 | 0.0185 | 125.28 | $< 2 \times 10^{-16}$ |
| Degree 1 (linear term) | 12.8359 | 0.4789 | 26.80 | $< 2 \times 10^{-16}$ |
| Degree 2 (quadratic term) | 6.9187 | 0.4789 | 14.45 | $< 2 \times 10^{-16}$ |
| Degree 3 (cubic term) | 4.0667 | 0.4789 | 8.49 | $< 2 \times 10^{-16}$ |
| Degree 4 (quartic term) | 2.1580 | 0.4789 | 4.51 | $7.81 \times 10^{-6}$ |

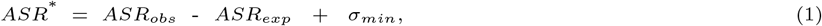

where *ASR*_*obs*_ represents the observed alternative splicing values computed from the annotated files, and *ASR*_*exp*_ corresponds to the expected values derived from the fitted polynomial model. The difference between the observed and expected values centers the data around zero, highlighting deviations as positive (above expected) or negative (below expected). However, values below 1 (*ASR*^*^ *<* 1) lack biological meaning. Thus, we added a normalization constant:

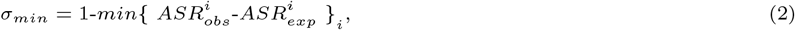

which adjusted all values so that the minimum normalized *ASR*^*^ is equal to 1. This correction resulted in a normalized measure, ASR^*∗*^, which preserves relative differences in alternative splicing while reducing biases introduced by annotation evidence. As a result, this metric allows for more accurate cross-species comparisons and provides a more reliable basis for downstream analyses of splicing complexity. As shown in Figure 2(C), and as expected after normalization, the correlation between ASR^*∗*^ and the percentage of fully supported CDSs becomes non-significant, with a coefficient close to zero. As a consequence, the normalization has effectively removed the bias introduced by the EGAP pipeline. However, it is important to emphasize that these normalized values should be interpreted as representative estimates rather than precise measurements, given that the normalization process is based on an empirical approach.

## Discussion

In this study, we systematically investigated how annotation quality affects estimates of alternative splicing levels across 670 multicellular eukaryotic species annotated by the NCBI Eukaryotic Genome Annotation Pipeline. We collected a total of 23 annotation-related variables, which are organized into three major groups: (1) assembly quality metrics, (2) evidence-based annotation indicators, and (3) annotation origin of CDSs. This categorization accounts for the methodological heterogeneity arising from the integration of diverse annotation strategies across species, capturing differences in genome sequencing, assembly contiguity, the amount and type of supporting evidence, and the extent to which coding sequences are derived from computational prediction versus experimental validation.

Although most annotation-related variables showed only weak or negligible correlations with alternative splicing levels, the percentage of fully supported CDSs emerged as the dominant predictor in both pairwise and multivariate analyses. Our findings imply that annotations heavily reliant on computational prediction without sufficient experimental validation may systematically underestimate splicing complexity. Conversely, species with a higher proportion of empirically supported CDSs exhibited elevated alternative splicing levels. As a consequence, rather than being directly driven by the volume of raw transcriptomic or proteomic data—as previously emphasized [Buen Abad Najar et al., 2020, Shen et al., 2014]—splicing estimates are more strongly influenced by how effectively this empirical evidence is incorporated into the annotation process. It may have important implications for comparative studies, as it suggests that cross-species differences in splicing diversity could partly reflect differences in annotation practices rather than true evolutionary divergence. This, in turn, reveals that the percentage of fully supported CDSs can serve as a key proxy for how much experimental evidence is actually reflected in isoform annotations, offering a practical handle to correct potential biases in splicing estimates.

Another important result is the apparent lack of association between assembly quality metrics and estimates of alternative splicing. While variables such as contig N50, scaffold count, and gap length showed only weak correlations with splicing levels, this finding must be interpreted with caution. Our dataset was specifically filtered to include only high-quality assemblies—those at the chromosome or complete genome level—which minimizes variability in assembly contiguity. As a result, potential biases linked to more fragmented assemblies may have been underrepresented. In studies that include lower-quality or more heterogeneous assemblies, such as those at the scaffold or contig level, assembly completeness has been shown to significantly influence gene and transcript annotation outcomes [Salzberg, 2019, Yandell and Ence, 2012]. Thus, poorly assembled genomes may fail to recover splice variants. Our findings further suggest that annotations with an excessive reliance on *Homo sapiens* data may lead to an overestimation of alternative splicing levels. This pattern suggests that homology-based annotation, while useful for transferring known gene models across species, may also introduce biases in isoform prediction by over-representing well-characterized transcript structures from model organisms [Steijger et al., 2013]. Such biases are especially significant in non-model species, where scarce species-specific data can misrepresent the true diversity of transcripts. Future work should prioritize integrating more experimental data, especially long-read RNA-sequencing. Unlike short-read methods, which often struggle to reconstruct complete transcript isoforms, long-read sequencing can directly capture full-length transcripts, providing an accurate and comprehensive view of transcript diversity [Wang et al., 2016, Weirather et al., 2017, Tardaguila et al., 2018]. Incorporating long-read datasets into genome annotations would substantially reduce biases introduced by incomplete or fragmented transcript reconstructions, and offer a more faithful representation of isoform complexity across species.

In summary, our results reveal that key features of the annotation process, such as the proportion of CDSs supported by empirical data, introduce systematic biases in estimates of splicing complexity. To correct for this annotation-driven bias, we introduced a normalization procedure based on a polynomial regression between ASR and the percentage of fully supported CDSs. The resulting adjusted metric, ASR*, effectively removed the correlation with annotation evidence, enabling more robust comparisons across species. Our normalized metric preserves relative differences in splicing while mitigating artifacts introduced by heterogeneous annotation methodologies. This correction strategy has already proven effective in a previous study [de la Fuente et al., 2025], where it enabled a large-scale comparative analysis of alternative splicing patterns across diverse species. Our work complements previous efforts to benchmark genome annotation accuracy [Frankish et al., 2018] and adds a new layer by quantifying its impact on transcriptomic metrics. As reference databases such as RefSeq and Ensembl continue to expand, and as long-read sequencing technologies become more accessible, integrating detailed metadata will be critical to disentangle biological signal from technical noise. Finally, adopting standardized evidence-weighted annotation practices and increasing transparency in annotation pipelines will improve the reproducibility and interpretability of cross-species analyses of alternative splicing.

## Supplementary Material

Supplementary Material is available at NAR Genomics and Bioinformatics Online.

## Data Availability

Availability of data and materials: Data was downloaded in November of 2024 from the NCBI database [Kitts et al., 2015]. The list of species under study and the datasets generated in this study are available in Zenodo, at https://doi.org/10.5281/zenodo.15274592.

## Author Contributions Statement

Conceptualization, R.d.l.F., W.D.-V., V.A. and A.M.; methodology, R.d.l.F.; writing, R.d.l.F.; review and editing, R.d.l.F., W.D.-V., V.A. and A.M.; supervision, A.M. All authors have read and agreed to the actual version of the manuscript.

## Conflict of Intetest Disclosure

No competing interest is declared.

