## Supplementary Material for "Evidence-driven biases in alternative splicing inferred from NCBI Eukaryotic Genome Annotation Pipeline metadata"

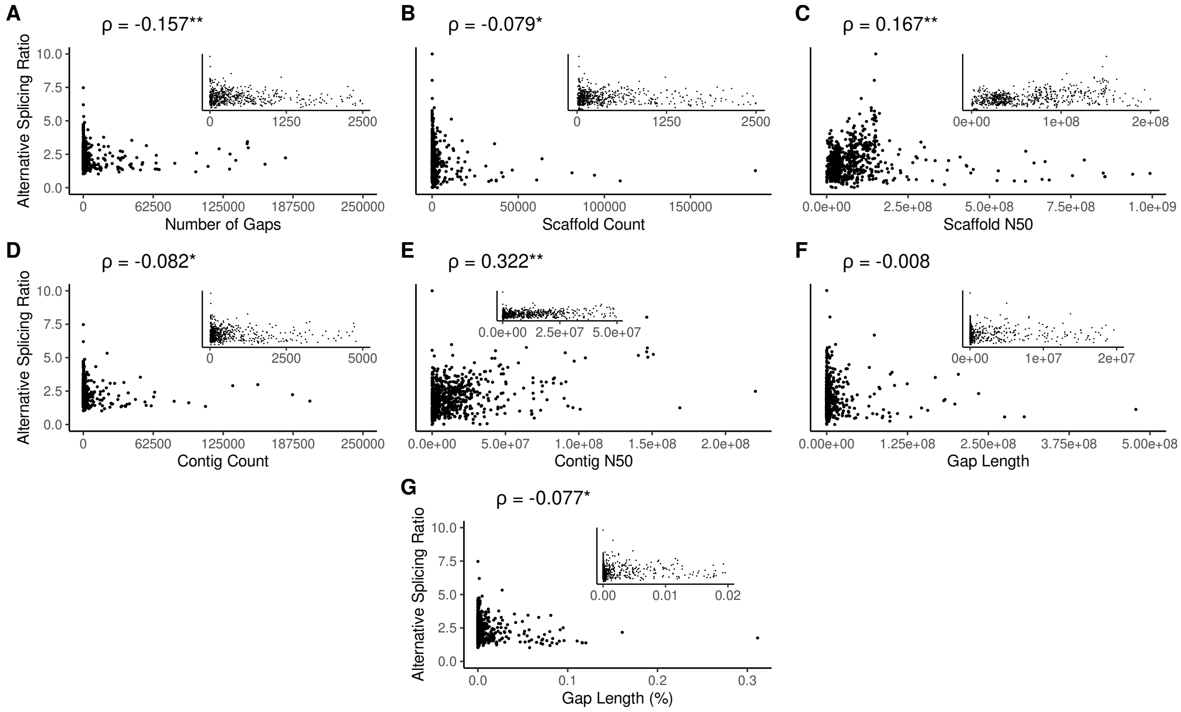


**Fig. 1.** Spearman correlation between the alternative splicing ratio and the genome assembly quality metrics: number of gaps, scaffold count, scaffold N50, contig count, contig N50, gap length, and gap length (%).

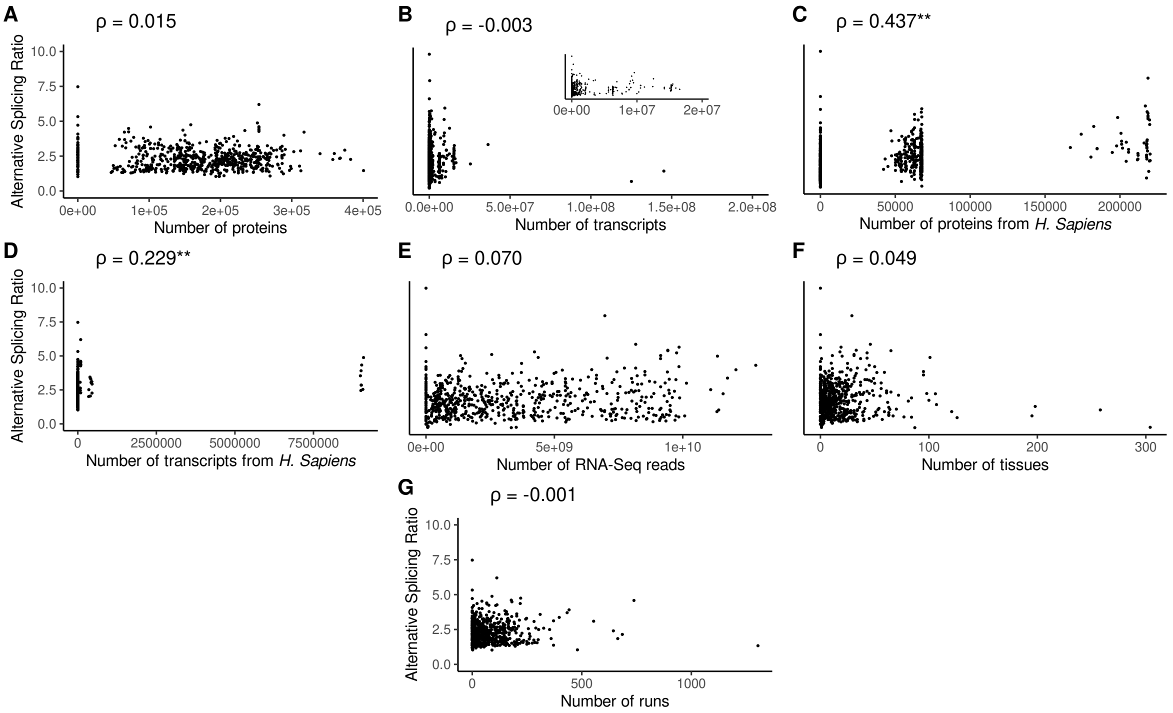


**Fig. 2.** Spearman correlation between the alternative splicing ratio and the experimental evidence metrics: number of proteins, number of transcripts, number of proteins from *Homo sapiens*, number of transcripts from *Homo sapiens*, number of RNA-Seq reads, number of tissues, and number of experimental runs.

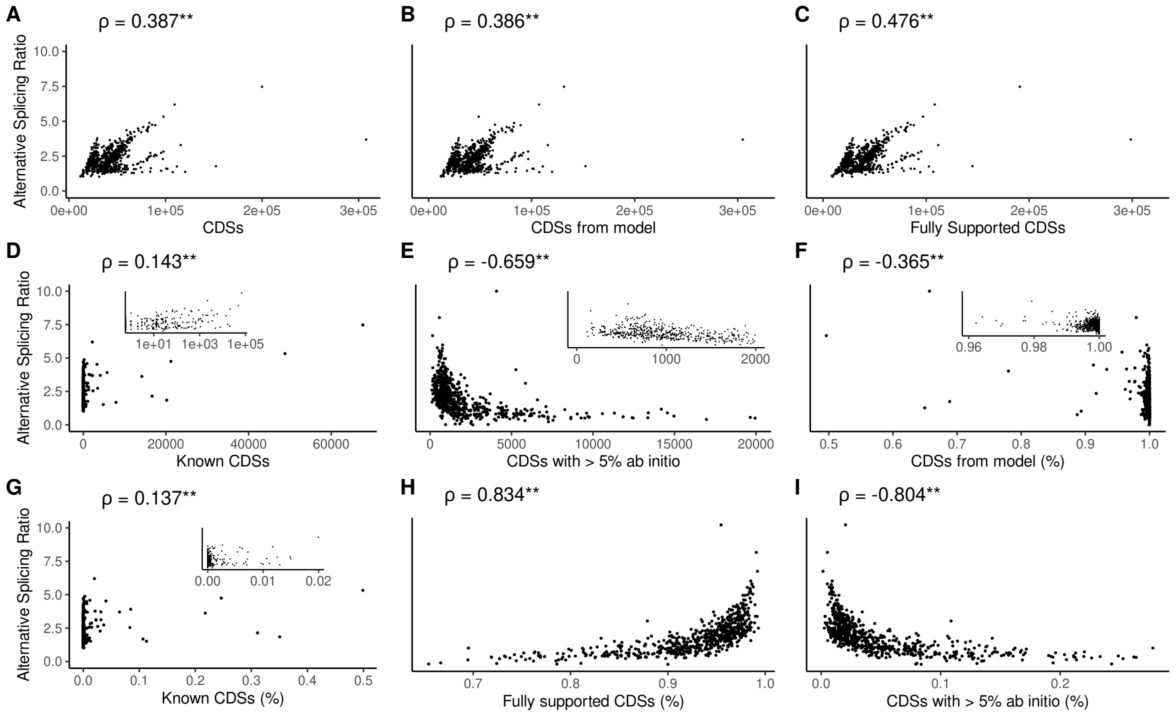


**Fig. 3.** Spearman correlation between the alternative splicing ratio and the CDS-related annotation metrics: total number of CDSs, number of CDSs from model, number of fully supported CDSs, number of known CDSs, number of CDSs for which more than 5% of their sequence is annotated using ab initio prediction, percentage of CDSs from model, percentage of known CDSs, percentage of fully supported CDSs, and percentage of CDSs for which more than 5% of their sequence is annotated using ab initio prediction.
